# A novel fermented Yi traditional medicine efficiently suppresses the replication of SARS-CoV-2 in vitro

**DOI:** 10.1101/2020.12.29.424534

**Authors:** Shisong Fang, Benhong Xu, Xiangrong Song, Wukun Liu, Yongmei Xie, Xifei Yang

**Author notes:** These authors contributed equally to this work (Shisong Fang, Benhong Xu). **Corresponding author:** E-mail addresses (Xifei Yang), (Yongmei Xie).

## Abstract

Currently, the coronavirus disease 2019 (COVID-19) caused by the severe acute respiratory syndrome coronavirus 2 (SARS-CoV-2) has a worldwide epidemic, causing more than 80 million infections and more than 1.7 million deaths. The pandemic has led to the closure of enterprises and schools in many countries, resulting in serious disruption of the global economy and social activities. Remdesivir is currently approved by the FDA for the treatment of COVID-19, but the WHO declared that Remdesivir is almost ineffective against COVID-19. The research and development of vaccines has made great progress, but it will take at least several months for safe and effective vaccines to be widely used clinically. Clinical studies revealed that some Traditional Chinese Medicines, such as Lianhua Qingwen Capsule and Huoxiang Zhengqi Water, exhibited excellent therapeutic effect on COVID-19. However, until now, there is still no cure for COVID-19. Therefore, there is an urgent need to find medicines that can effectively fight against the SARS-CoV-2. In this study, JIE BEN No. 10 (JB10), a fermentation broth produced by Yi traditional medicine fermentation technology, was explored for its anti-coronavirus activity. The *in vitro* data showed that JB10 could significantly suppresses the replication of the SARS-CoV-2 with an EC_50_ of 769.1 times dilution and a selection index of 42.68. Further studies indicated that JB10 had significant anti-inflammatory and antioxidant activities. The analysis of active components suggested that JB10 contained a large amount of superoxide dismutase (SOD), flavones, polyphenols, crude polysaccharide, etc. which may explain the anti-coronavirus activity, anti-inflammatory and antioxidant effects. Our study provides a new potentially therapeutic strategy for COVID-19.

## Introduction

Coronavirus disease 2019 (COVID-19) is an acute respiratory infection caused by SARS-CoV-2. The main symptoms of patients are fever, fatigue, tussiculation, rhinobyon, runny nose, etc. About half of the patients will have difficulty breathing within a week, and severe cases will rapidly progress to acute respiratory distress syndrome, septic shock, difficult to correct metabolic acidosis and coagulopathy. ^1^ The SARS-CoV-2 belongs to the β genus of coronaviruses. It has an envelope and the particles are round or oval, often pleomorphic, with a diameter of 60-140nm. ^2^ At present, the SARS-CoV-2 has caused more than 80 million infections and more than million deaths in the worldwide according to Johns Hopkins University data. The epidemic has caused many countries to close enterprises and schools, causing severe damage to the global economy. At present, Remdesivir has been approved by the FDA for the treatment of patients with COVID-19, ^3^ but the WHO claims that Redesivir is almost ineffective against COVID-19. ^4^ The research and development of vaccines has made great progress, but safe and effective vaccines are widely used in clinical practice, and it will take at least several months.

At present, the COVID-19 epidemic has been well controlled in China. In this process, traditional Chinese medicine has made great contributions. For example, Lianhua Qingwen granules and Huoxiang Zhengqi dropping pills can significantly improve the symptoms of fever, cough, sputum and breathing difficulties caused by the novel coronavirus. ^5^ However, there is still no specific medicine to treat the COVID-19. Therefore, the discovery of medicines that can significantly inhibit the replication of the novel coronavirus is vital to the prevention and control of the COVID-19 epidemic.

Yi traditional medicine is an important part of Traditional Chinese Medicine (TCM) and has made great contributions to the development of Chinese civilization. Among them, the world-famous Yunnan Baiyao is derived from the ancient Yi prescription. ^6^ JB10, a fermentation broth, is fermented from Phyllanthus emblica, Rosa roxburghii Tratt., Lemon, Gastrodia elata Bl. and honey according to Yi traditional medical theory. ^7^ In this work, in the process of screening anti-SARS-CoV-2 candidate drugs, we found that JB10 can significantly inhibit the replication of the novel coronavirus. Further research shows that JB10 has excellent antioxidant and anti-inflammatory effects. This research will provide new methods for anti-novel coronavirus treatment.

## Results

### *In vitro* antiviral activity against SARS-CoV-2 replication

Standard assays were carried out to measure the cytotoxicity effects of JB10. The cytotoxicity of JB10 in Vero E6 cells was determined by the CCK8 assay. As can be seen from Figure 1, JB10 did not show significant toxicity on Vero E6 cells at concentrations above 100 times diluted concentration. The CC_50_ was 18.02 times diluted concentration. Thus, the least dilution time for JB10 was set as 100 times. The Vero E6 cells were infected according to Wang et al. ^8^ with some modofications with SARS-CoV-2 at a multiplicity of infection (MOI) of 0.05 in the presence of varying concentrations of the candidate drug (diluted to 100, 200, 400, 800, 1600 and 3200 times). The viral replication was evaluated by quantification of viral copy numbers in the cell supernatant by quantitative real-time RT-PCR (qRT-PCR). As shown in Figure 1, JB10 exhibited potent antiviral against the SARS-CoV-2 (EC_50_=769.1 times dilution). The selective index is 42.68. These results indicated that JB10 may be developed as a candidate for COVID-19 treatment. Further in vivo evaluation of this candidate against SARS-CoV-2 infection is recommended.

**Figure 1.**
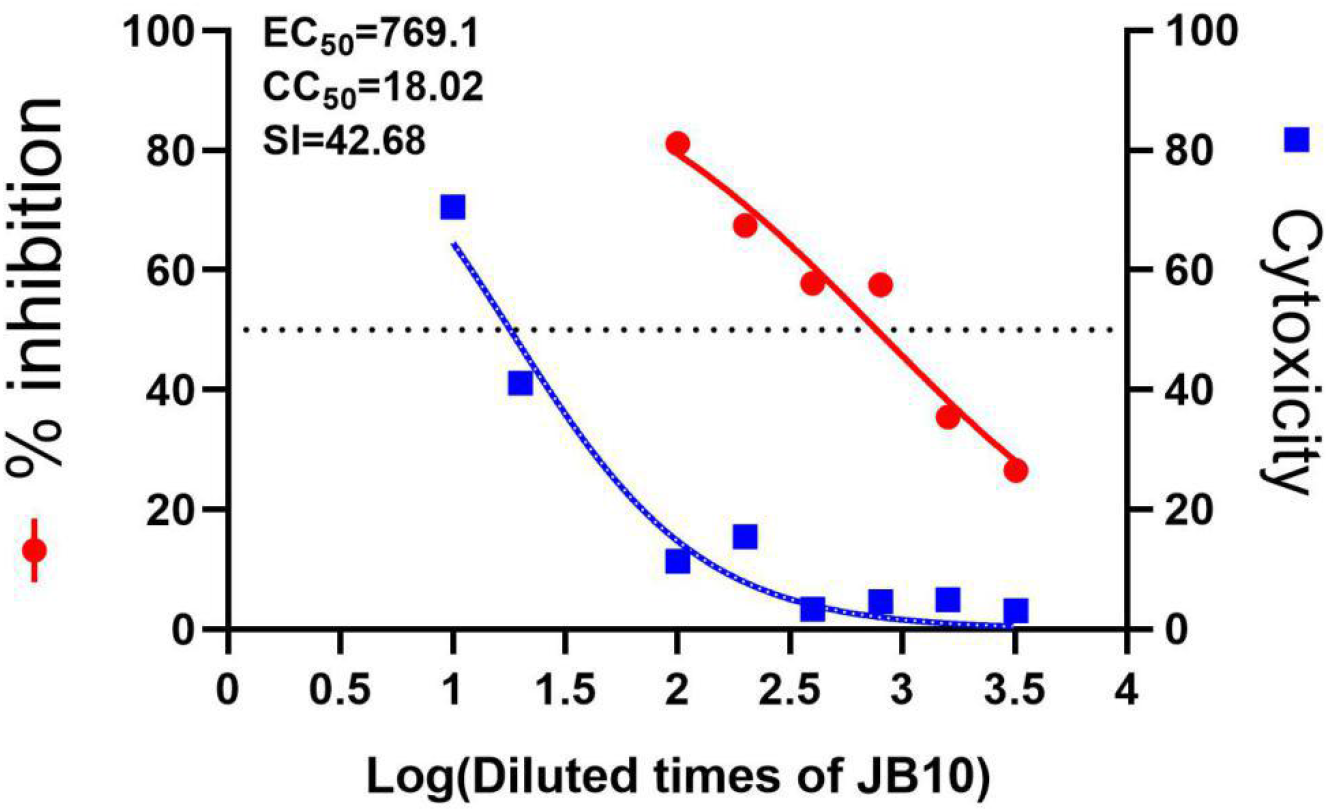
The antiviral activities of JB10 against SARS-CoV-2 *in vitro*. The Vero E6 cells were infected with SARS-CoV-2 at an MOI of 0.05 in the presence of different doses of No.10 fermentation liquor for 24 h. The viral yield in the cell supernatant was then quantified by qRT-PCR. Cytotoxicity of No.10 fermentation liquor to Vero E6 cells was measured by CCK-8 assays. The experiments were repeated at least 3 times.

### Antioxidant activities

Oxidative stress, characterized by excess levels of reactive oxygen species (ROS), plays a vital role in the progression of COVID-19.^9^ Many evidences support therapeutic counterbalancing of ROS by antioxidants can prevent COVID-19 from becoming severe. ^10,11^ In order to evaluate the antioxidant activities of JB10, DPPH free radical and hydroxyl radical scavenging assay were performed. As shown in Figure 2A, JB10 exhibited very strong and significant dose-dependent DPPH radical scavenging activities. When JB10 was diluted to 800, 1600, 6400, and 12800 times, the DPPH free radicals were removed 82.99%, 65.23%, 53.09% and 25.84%, respectively. As can be seen from Figure 2B, JB10 also displayed strong hydroxyl radical scavenging activities. After diluted to 20, 30, 40, and 50 times, the hydroxyl radicals were removed 67.42%, 59.92%, 53.91% and 51.64%, respectively.

**Figure 2.**
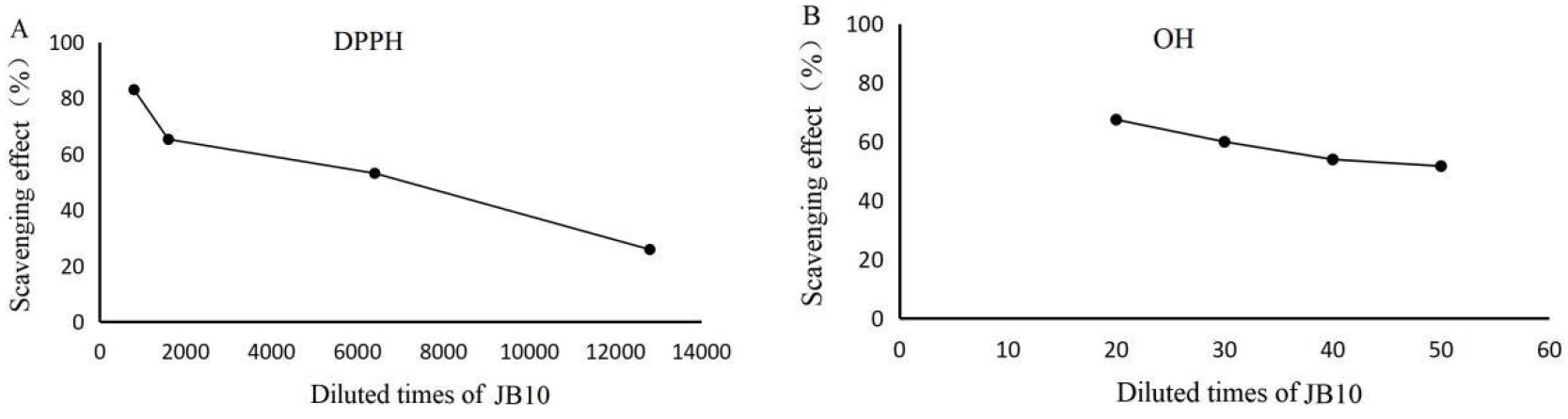
Effect of JB10 on DPPH radical scavenging activity and hydroxyl radical scavenging activity. A. The JB10 was diluted for 800, 1600, 6400, and 12800 times. B. The JB10 was diluted for 20, 30, 40, and 50 times.

### Anti-inflammatory activity

Pulmonary hyper-inflammation and potentially life-threatening “cytokine storms” are the characteristics of COVID-19, which cause disease severity and death in infected patients. ^12^ According to the publication, controlling the local and systemic inflammatory response in COVID-19 may be as important as anti-viral therapies. ^13^ The anti-inflammatory activity was determined according to Liu ^14^ with some modifications. Before starting the experiment of anti-inflammatory activity, the cytotoxicity of JB10 to RAW 264.7 cell line was tested for 24 h by MTT assay. As shown in Figure 3A, there was no significant toxicity up to diluted 160 times on RAW 264.7 cells.

**Figure 3.**
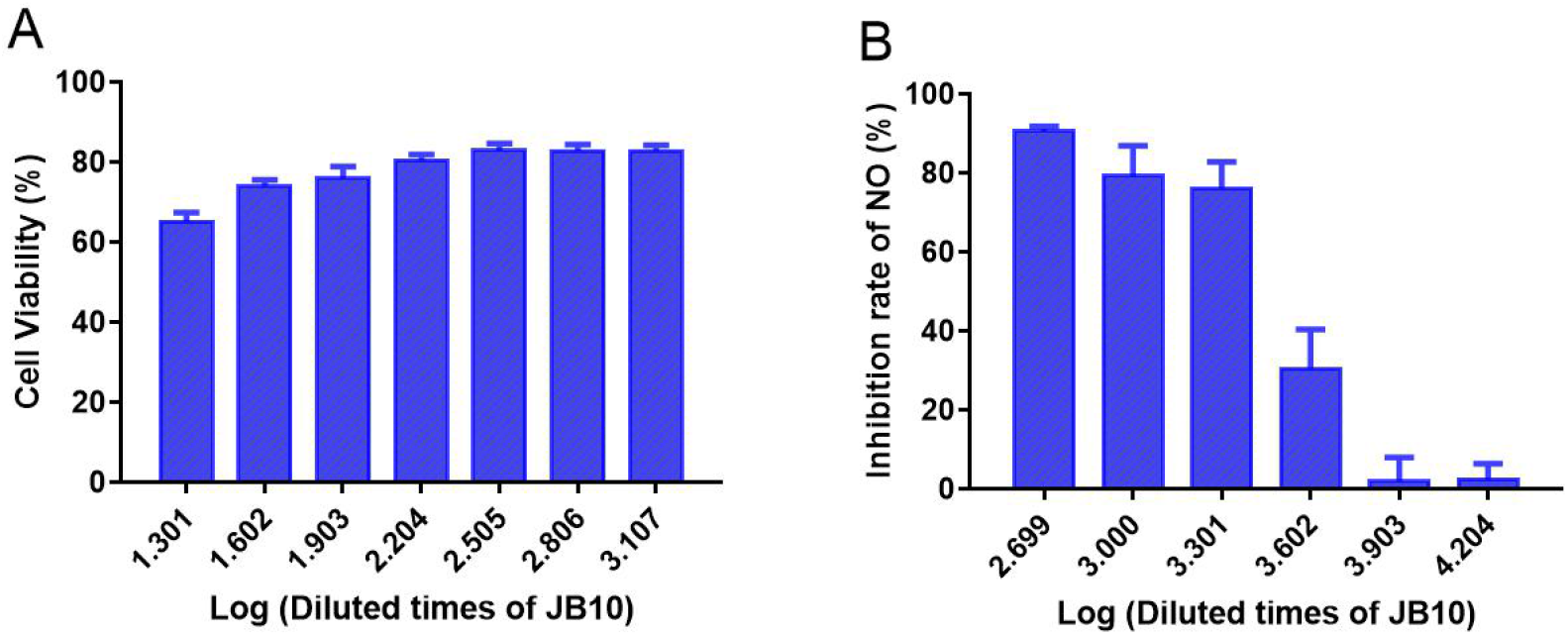
Anti-inflammatory activity of JB10 A. The cytotoxicity of JB10 to RAW 264.7 cell line; B. Inhibition of NO production.

The anti-inflammatory activity of JB10 was evaluated in LPS-induced RAW264.7 cells under the following concentrations (diluted to 500, 1000, 2000, 4000, 8000, and 16000 times). As can be seen from Figure 3B, JB10 could effectively suppress the NO production. When the JB10 was diluted to 2000 times, the inhibition rate of NO rate was still close to 80%. These results suggested that JB10 has potent activity against inflammation.

### Analysis of active components

Since JB10 has potent antiviral activities against SARS-CoV-2, antioxidant activities and anti-inflammatory activity, we analyzed its components to provide evidences for JB10. As can be seen from Table 1, JB10 contains a lot of active components including Vitamin C, SOD, flavones, polyphenols, crude polysaccharide, proanthocyanidins, saponins and coumarin etc. These substances have antioxidant, anti-inflammatory, immune and antiviral effects.

**Table 1.** Active components in JB10

| Components | Content |
| --- | --- |
| SOD | 2398.4 U/g |
| Vitamin C | 550 mg/100ml |
| Flavones | 893.17 mg/100ml |
| Polyphenols, | 1243.50 mg/100ml |
| Crude polysaccharide | 1753.49 mg/100ml |
| Proanthocyanidins | 303.17 mg/100ml |
| Saponins | 330.38 mg/100ml |
| Coumarin | 49.50 mg/100ml |

## Discussion

TCM has a nearly 3000 years of history and is widely practiced in China. Several epidemics have occurred in the history of China. But, fortunately, no large numbers of casualties occurred, in part due to the use of TCM. ^15^ Yi traditional medicine is a vital part of TCM and has its unique medical theory and practical experience. ^16^ JB10 is a fermentation broth from Phyllanthus emblica, Rosa roxburghii Tratt, Lemon, Gastrodia elata Bl. and honey according to Yi traditional medical theory. In this work, the antiviral activity against the SARS CoV-2 was investigated. We observed significant, potent, and dose-dependent antiviral activity of JB10. As expected from the fact that Yi traditional medicine was widely used for long time, the treatment did not exert cytotoxicity. Oxidative stress and inflammation play important roles in the progression of COVID-19. We also explored the anti-inflammatory and antioxidant activities of JB10. The results showed that JB10 has potent anti-inflammatory and antioxidant activities.

JB10 contains a lot of Vitamin C, SOD, flavones, polyphenols, crude polysaccharide, proanthocyanidins, saponins and coumarin etc. Vitamin C has multiple pharmacological characteristics, antiviral, anti-oxidant, anti-inflammatory and immunomodulatory effects, which make it a potential therapeutic option in management of COVID-19. ^17^ SOD, one of the powerful antioxidants, plays a major role in fighting against free radical damage and inflammation, and SOD might exert an anti-aging effect. While, aging immunity may exacerbate COVID-19. ^18^ So, SOD may be beneficial for the treatment of COVID-19. A previous study showed that flavonoids could inhibit key proteins in the coronavirus infection cycle and SARS-CoV proteases PLpro and 3CLpro. ^19^ Flavonoids can also inhibit NTPase/helicase and N protein of SARS-CoV. ^20^ JB10 contains high levels of total flavone, which might be involved in the inhibitory effects of the virus replication as we observed. Polyphenols, a major class of bioactive compounds in nature, are known for their antiviral activity and pleiotropic effects. In a previous report, polyphenols was shown to inhibit SARS-CoV-2 fusion/entry, disrupt SARS-CoV-2 replication and suppress the host inflammatory response. ^21^ From Table 1, the content of polyphenols in JB10 is abundant. This may also contribute to the antiviral activity of JB10. Polysaccharides have broad applications in anti-virus, especially in anti-coronavirus due to the good safety, immune regulation and antiviral activity. ^22^ Song et al. found that polysaccharides exhibited potent inhibitory activity against SARS-CoV-2. A further study suggested that polysaccharides could bind to the S glycoprotein to prevent SARS-CoV-2 host cell entry. ^23^ Zhu et al. used main protease (M^pro^) of SARS-Cov-2 to screen plant flavan-3-ols and proanthocyanidins. The results showed that proanthocyanidins could inhibit the M^pro^ activity of SARS-Cov-2. ^24^ The high levels of proanthocyanidins in the JB10 might also contribute to the inhibitory effects of SARS-Cov-2. Recent modeling studies have shown that the saponin derivative has a high binding affinity to the papain-like protease of SARS-CoV-2 ^25^. Molecular docking analysis also showed that the natural coumarin analog toddacoumaquinone had a significant inhibitory ability on the main protease of COVID-19 (compared to the complex α-ketoamide (PDBID: 5N5O)), with a binding energy of −7.8kcal/ mol. The synthesized coumarin analog (1m) also showed considerable inhibitory ability, and the binding energy to the main protease of COVID-19 (containing α-ketoamide) was −7.1kcal/mol. JB10 contains high amount of coumarin, which might contribute to the inhibitory effects of the virus replication as we observed. ^26^

In summary, given the urgency of the pandemic around the world, such safe and cheap liquor might provide help for prevention or treatment of COVID-19 patients. Our data support that the JB10 goes into the clinical stage for further validation of the efficiency for anti-SARS-CoV-2 infection and therapy.

## Materials and Methods

### Materials

Vero E6 cell line and murine macrophage RAW 264.7 cell line were purchased from ATCC (Manassas, VA, USA). The Vero E6 cell line was cultivated in high glucose 278 Dulbecco`s minimal essential medium (Gibco 41966-029) supplemented with 10% (v/v) FBS, penicillin, and streptomycin at 37 °C in an atmosphere of 5% CO_2_. The RAW 264.7 cell line was cultured in Dulbecco’s modified eagle’s medium (American Type Culture Collection) supplemented with 10% heat-inactivated fetal serum, and streptomycin, and penicillin. The 281 SARS-CoV-2 strains were obtained from Shenzhen Center for Disease Control and Prevention. Viral titers were determined by TCID_50_ titration. JIE BEN No.10 fermentation broth was supplied by Guizhou Jinqianguo Biotechnology Co. Ltd. Lipopolysaccharides (LPS), MTT, Ferrous Sulfate and 2,2-diphenyl-1-picrylhydrazyl were obtained from Sigma-Aldrich (St. Louis, MO, USA). Salicylic acid was purchased from J&K Scientific Ltd. (Beijing, China). Nitric oxide (NO) assay kit (Cat. No.: S0021S) was obtained from Shanghai Beyotime Biotechnology Co. Ltd. (Shanghai, China).

### *In vitro* antiviral activity

The Vero E6 cells were plated into 96-well plates at densities of 1×10^4^ cells/well. The cells were infected with SARS-CoV-2 at an MOI of 0.05 for 2 h, then the virus containing medium was replaced with fresh medium in the presence of different doses of JB10 for 24 h. The viral yield in the cell supernatant was then quantified by qRT-PCR. Cytotoxicity of JB10 to Vero E6 cells was measured by Cell Counting Kit-8 (CCK8) assays as previously reported. Cells were seeded into 96-well plates (5×10^3^ cells/well) and cultured overnight. JB10 was diluted to 10, 20, 100, 200, 400, 800, 1600 and 3200 times with fresh medium and incubated with cells for another 48 h, the CCK8 was added and incubated for another 4 h. The absorbance at 450 nm was measured with a microplate reader. The cytotoxicity and mean % inhibition of virus yield of JB10 were shown in Figure 1. The experiments were repeated at least 3 times.

### Antioxidant activities

The antioxidant activities of JB10 were evaluated by DPPH free radical scavenging assay and hydroxyl radical scavenging assay according to the slightly modified method by Lim et al. ^27^ JB10 was diluted with distilled water for different concentrations of sample. For DPPH free radical scavenging assay, two hundred microliters sample and 2 ml 400 μL DPPH solution (dissolved in 70% ethanol solution) were reacted at room temperature for 1 h. The absorbance was measured at 517 nm with a blank containing DPPH and 70% ethanol solution. The DPPH radical scavenging rate (%) was calculated using the following equation:

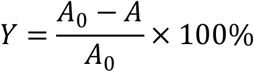

Where, A_0_ is the absorbance value of the blank control, A is the absorbance value of the samples.

For hydroxyl radical scavenging activity assay, 250 μL FeSO_4_ solution (2 mM), 250 μL salicylic acid solution (2 mM) and 100 μL sample were mixed well. Two hundred and fifty microliters hydrogen peroxide solution (0.01%) was added and the reaction mixture was stirred for 30 min at 37 °C. The absorbance was determined at 510 nm after cooling. The hydroxyl radical scavenging rate (%) was calculated according to the following equation:

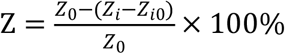

Where, Z_0_ is the absorbance value of the blank control, Z_i_ is the absorbance value of the sample, Z_i0_ is the absorbance value of the H_2_O_2_ without the color agent.

### Anti-inflammatory activity

Before starting the experiment of anti-inflammatory activity, the cytotoxicity of JB10 to RAW 264.7 cell line was tested. RAW 264.7 cells were seeded into 96-well plates. After overnight incubation, the culture medium was replaced by fresh medium containing gradient concentrations (diluted to 0, 20, 40, 80, 160, 320, 640 and 1280 times) of JB10. Triplicate wells were tested at each concentration. After 24 h treatment, the cell viability was determined by MTT assay. The anti-inflammatory activity of JB10 was performed according to Liu et al. ^14^ with some modifications. Briefly, RAW 264.7 cells were incubated in 24-well plates with the density of 100, 000 cells/well. After overnight incubation, the culture medium was replaced by 0.5 ml fresh medium containing different concentrations of JB10. Triplicate wells were tested at each concentration. After 1 h incubation, 50 μL medium containing LPS was added into each well. The final concentration of LPS was 1μg/mL. After 18 h culture, 100 μL supernatant was put into another clean 96-well plates and the absorbance of the samples was then measured at 540 nm. The content of NO was calculated from a sodium nitrite standard curve. The inhibition rate of NO production was calculated as inhibition rate (%) = 100×(C_OD_ − S_OD_)/C_OD_. Where, C_OD_ is the control optical density, S_OD_ is the sample optical density. The IC_50_ value was defined as the concentration at which the nitrite radicals were reduced by 50%.

### Analysis of active components

The active components were determined by China National Research Institute of Food & Fermentation Industries (Beijing, China) according to relevant standards.

## Acknowledgments

This work was financially supported by the Shenzhen Bay Laboratory Opening Fund (SZBL202002271005), the Sichuan Province Science and Technology Plan for Science and Technology Department of Sichuan Province (2020YFS0008) and the Open Project of Chinese Materia Medica First-Class Discipline of Nanjing University of Chinese Medicine (No. 2020YLXK010).

